# Multi-task deep latent spaces for cancer survival and drug sensitivity prediction

**DOI:** 10.1101/2024.03.18.585492

**Authors:** Teemu J. Rintala, Francesco Napolitano, Vittorio Fortino

## Abstract

**Motivation:** Cancer is a very heterogeneous disease that can be difficult to treat without addressing the specific mechanisms driving tumour progression in a given patient. High-throughput screening and sequencing data from cancer cell-lines has driven many developments in drug development, however, there are important aspects crucial to precision medicine that are often overlooked, namely the inherent differences between tumours in patients and the cell-lines used to model them *in vitro*. Recent developments in transfer learning methods for patient and cell-line data have shown progress in translating results from cell-lines to individual patients *in silico*. However, transfer learning can be forceful and there is a risk that clinically relevant patterns in the omics profiles of patients are lost in the process.

**Results:** We present MODAE, a novel deep learning algorithm to integrate omics profiles from cell-lines and patients for the purposes of exploring precision medicine opportunities. MODAE implements patient survival prediction as an additional task in a drug-sensitivity transfer learning schema and aims to balance autoencoding, domain adaptation, drugsensitivity prediction, and survival prediction objectives in order to better preserve the heterogeneity between patients that is relevant to survival. While burdened with these additional tasks, MODAE performed on par with baseline survival models, but struggled in the drug-sensitivity prediction task. Nevertheless, these preliminary results were promising and show that MODAE provides a novel AI-based method for prioritizing drug treatments for high-risk patients.

**Availability:** https://github.com/UEFBiomedicalInformaticsLab/MODAE

## Introduction

High-throughput sequencing of omics data can be used to characterize the molecular state of cells and tissues in both patient tumors and cancer cell-lines that are used to model them. Very large patient cohorts such as the cancer genome atlas (TCGA) have been used to study the molecular mechanisms underlying cancer progression and the remarkable heterogeneity of tumors, leading to the development of precision medicine approaches that target specific genomic aberrations to improve treatment outcomes (Chin et al., 2011). On the other hand, cancer cell-lines have been widely screened *in vitro* for sensitivity with large panels of anticancer compounds and thoroughly characterized in projects such as CCLE (Ghandi et al., 2019; Barretina et al., 2012), GDSC (Yang et al., 2013), and CTRP (Seashore-Ludlow et al., 2015; Basu et al., 2013). Large collections of screening results together with comprehensive molecular profiles of the cell-lines provide incredible resources for data-mining drug mechanisms and targeted therapies. However, the heterogeneity of cancer aberrations and the intrinsic differences between cancer cell lines and patient tumors has made it difficult to identify appropriate models for specific (sub-)types of cancer (Salvadores et al., 2020). In order to translate drug screening results to patient tumors it is very important to properly account for these differences (e.g. by matching each patient tumour to a cell-line model based on the similarity of their omics profiles). However, instead of directly establishing the best representative model for each cancer, the issue could be addressed as a domain adaptation problem to translate predictive models learned on cell-line omics profiles to the domain of patient tumor omics profiles. Recently, deep learning approaches have been shown to outperform traditional machine learning and domain adaptation techniques in this complex task (He et al., 2022). Accurately translated sensitivity screens could help prioritize drugs to a given group of omics profiles and reduce drug development costs for precision oncology.

To address the problem of transfer learning, Dincer et al. (2020) implemented an adversarial extension of the autoencoder model (AD-AE) to obtain a representation of data free from confounding factors between datasets. CODE-AE-ADV by He et al. (2022) used a more robust version of the adversarial autoencoder model by utilizing the Wasserstein distance as inspired by WGAN (Arjovsky et al., 2017; Gulrajani et al., 2017). CODE-AE-ADV also implemented shared and private embeddings for learning separate latent representations of the confounders in addition to the deconfounded representation shared between data domains. The goal of CODE-AE-ADV was to obtain a representation that contains only the shared characteristics of both cell-line and patient tumor omics profiles and to use this representation to learn a transferable model of drug sensitivity prediction. In addition to bulk omics, a similar transfer learning schema has also been applied to single-cell drug sensitivity by Chen et al. (2022). However, focusing solely on optimizing drug sensitivity may skew the shared embedding space towards features more informative to cancer cell lines than to patient cases. For a comprehensive cancer precision medicine approach, it is essential that the genomic embedding space that connects cell-line and patient tumor omics profiles, remains relevant to cancer patients. This can be achieved by incorporating additional learning tasks focusing on clinical aspects, such as survival data or molecular subtypes. This ensures the genomic data space accurately reflects patient information as well as cancer cell line information, potentially facilitating the identification of clinically actionable cancer subtypes. By studying patients with similar genomic profiles and drug sensitivities, and linking these to specific survival outcomes, we can attain a true precision medicine approach. To integrate these disparate objectives within machine learning we can use multi-task learning (MTL) approaches. MTL can be used to learn multiple tasks simultaneously instead of building separate models for each task or learning in stages. The combination of autoencoders for feature extraction and predictive models as part of MTL has been previously utilized for drug-sensitivity prediction (Chen et al., 2022) as well as survival prediction (Vo et al., 2021) separately in the context of cancer omics. In this paper we implement a novel end-to-end deep learning architecture extending the CODE-AE-ADV to include patient survival prediction and a joint learning scheme to learn a latent space that is suitable for prioritizing drugs for groups of patients based on their omics profiles.

## Methods

### MODAE

#### Autoencoder

The autoencoder consists of an encoder module *E* : ℝ^*d*^ → ℝ^*p*^ and a decoder module *D* : ℝ^*p*^ → ℝ^*d*^. We denote the input omics with **x**, the bottle-neck layer as **z** = *E*(**x**) and the reconstructed omics as 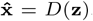. The purpose of the autoencoder is to find a low dimensional representation **z** ∈ ℝ^*p*^ of the data in an

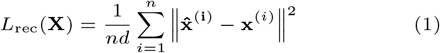

The input dimension *d* is included in the denominator in order to normalize the scales of different objectives for the multi-task learning step (Equation 8).

#### Adversarial deconfounding autoencoder

To separate learn separate representations for confounders, we split the vector **z** into three parts: **z**_*s*_ (shared), **z**_*p*_ (patient tissue), and **z**_*c*_ (cell-line). In contrast to He et al. (2022), we use a single encoder to learn all these representations by setting parts of **z** to **0** by applying the following function *f* before the decoder (i.e.,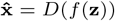):

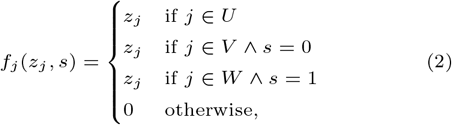

where *U* is the set of indices used for the shared representation, *V* is the set of indices for patients, and *W* is the same for cell-lines, and where *s ∈ {*0, 1*}* is an indicator for whether a given sample is from a cell-line (and not from patient tissue). Supervised tasks use the shared part and we denote *q* = |*U* | as its dimension. An adversary aims to distinguish cell-lines from patients and the WGAN inspired autoencoder is forced to learn representations that minimize differences between the two data domains, which should lead to more reliable transfer of predictions. The adversary and the autoencoder are trained in alternating steps. In the first step the critic network *C* : ℝ^*q*^ → ℝis trained to minimize the adversarial loss

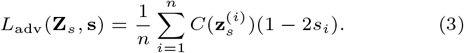

Then, the autoencoder is trained to minimize the sum of *L*_rec_ and the deconfounding loss

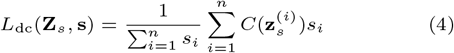

which is equivalent to the cell-line specific term of *L*_adv_ with the sign reversed. *L*_adv_ is an approximation of the Wasserstein distance between the distributions of the two sets of points in the latent representation (Arjovsky et al., 2017). In simpler terms, the critic or dataset detector network *C* is trained to estimate the patient-likeness of a given sample while the deconfounding autoencoder tries to minimize the distance of the cell-line sample distribution from the patient sample distribution. Furthermore, in order to satisfy the continuity assumptions of the WGAN model, we use a gradient penalty on *∇C*(**z**_s_) similarly to He et al. (2022). However, our formulation of *L*_adv_ and *L*_bc_ differs from the equivalent ℒ_critic_ and ℒ_gen_ in that we use the shared representation **z**_*s*_ while He et al. (2022) use a concatenation of shared and private representations and in our case the deconfounding loss is calculated on cell-lines instead of patient tissues.

#### Deep Cox regression for survival modeling

A deep learning extension of the non-parametric Cox regression model can be obtained with the the hazard function:

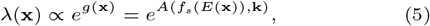

where *A* : ℝ^*q*^ → ℝis a survival risk prediction network, *f*_*s*_ : ℝ^*p*^ → ℝ^*q*^ is a function that extracts the shared representation from **z**, and **k** is an optional vector of additional input covariates, such as patient age or sex. We used Efron’s method (Efron, 1977) to account for tied survival times which can be quite common for large datasets or aggressive cancers. Hence, the loss function for training *A* is based on the adjusted Cox partial log-likelihood

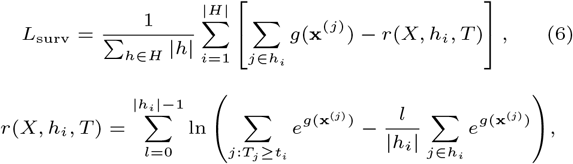

where *h*_*i*_ *∈ H* is the set of indices of all events for which *T*_*j*_ = *t*_*i*_∀*j* ∈ *h*_*i*_, *H* contains the sets of events for each unique time point *t*_*i*_, and **x**^(*i*)^ is the vector of omics data for patient *i*. To improve the numerical stability of the calculations we utilized the log-sum-exp trick. The likelihood was divided by the number of observed events to maintain consistent scaling of the losses.

#### Deep regression for drug sensitivity modeling

Drug sensitivity prediction was addressed as a multiple output regression problem. Since every drug-cell-line pair was not screened in CTRP, losses were computed on non-missing values only. In other words we minimize the loss

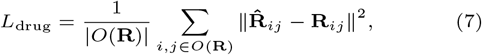

where **R** is a matrix of observed drug data that may contain missing values, *O* is a function that returns the set of indices of observed cell-line drug pairs, and 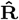 contains drug-sensitivity vectors 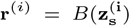 computed for each cell-line *i* with the drug-sensitivity network *B* : ℝ^*q*^ → ℝ^*m*^, where *m* is the number of drugs.

#### Joint multi-task learning

To simultaneously learn to represent the molecular state, to find a shared deconfounded latent space, and to predict both patient survival and cell-line drug sensitivity from omics data we define the total loss as a weighted sum of the losses used for individual tasks as follows:

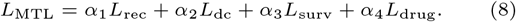

The weights *α*_1*−*4_ can be used to adjust the task priority and were tuned as hyper-parameters to explore different trade-offs as described in section 3.2.

To ensure robust training for the multi-task model, we implemented a pre-training procedure for each network module that corresponds to a specific task (Supplementary Algorithm 1). The autoencoder, critic network, deconfounding autoencoder, patient-survival network, and drug-sensitivity network can be trained in order before running the joint learning that updates the full model at each step (Supplementary Algorithm 2). In practice, skipping the first two steps and starting from the deconfounding autoencoder training worked better than pre-training each step. Pre-training of the deconfounding autoencoder was continued for 300 epochs, followed by 100 epochs each of patient survival and cell-line drug-sensitivity module training, after which the joint multitask training was run for 300 epochs. Algorithm 1 shows the joint learning procedure which samples equally sized minibatches from both datasets and updates the weights of all MODAE modules based on Equation 8. Since the number of patients and cell-lines can be different, we employed an oversampling strategy to generate the same number of minibatches from both sample types for each epoch. Supplementary Figure S1 shows the constituent losses for each epoch across training phases and cross-validation folds for a model selected by parameter search. We used the scaled exponential linear units (Klambauer et al., 2017) to increase the robustness of the joint model in place of more conventional batch-normalization. Furthermore, L2-norm regularization was applied on all the model weights, and dropout was applied after all layers except outputs and bottle-neck. The critic network *C* was trained in epochs that alternate with the autoencoder epoch in pretraining and multi-task epoch in joint training. Additionally, for each epoch of the deconfounding and joint training the adversary was trained for 5 epochs and used the Adam optimizer with recommended momentum parameters *β*_1_ = 0, *β*_2_ = 0.9 as suggested by Arjovsky et al. (2017); Gulrajani et al. (2017).

### Data and pre-processing

To perform a case-study on real patient data, we used cancer tissue data from the Swedish SCANB breast cancer cohort (Staaf et al., 2022) as a discovery dataset and the TCGA BRCA cohort as a validation dataset. To simplify the analysis, we focused on RNA-Seq data since the transcriptome has been shown to be the most relevant omic for drug sensitivity prediction (Costello et al., 2014). The SCANB datat was already pre-processed to fragments per kilobase per million (FPKM) while pre-processed TCGA data downloaded with the *curatedTCGAData* R-package (Ramos et al., 2020) was in upper-quartile (UQ) normalized transcripts per million (TPM). To harmonize them, SCANB data was normalized to UQ TPM using the *edgeR* R-package and then both were log-transformed to log_2_(TPM + 1). Cell-line transcriptome profiles were acquired from the DepMap portal (Corsello et al., 2020) which were already equivalently processed. Genes with average log-expression less than 1.0 in either the patient and cell-line dataset were excluded and the values in each dataset were independently centered and scaled to mean 0 and standard deviation 1. This yielded gene-expression datasets of 12021 genes, 9206 patients for discovery, 1093 patients for validation, and 1293 cell-lines for discovery.

#### Algorithm 1

Multi-task training of MODAE

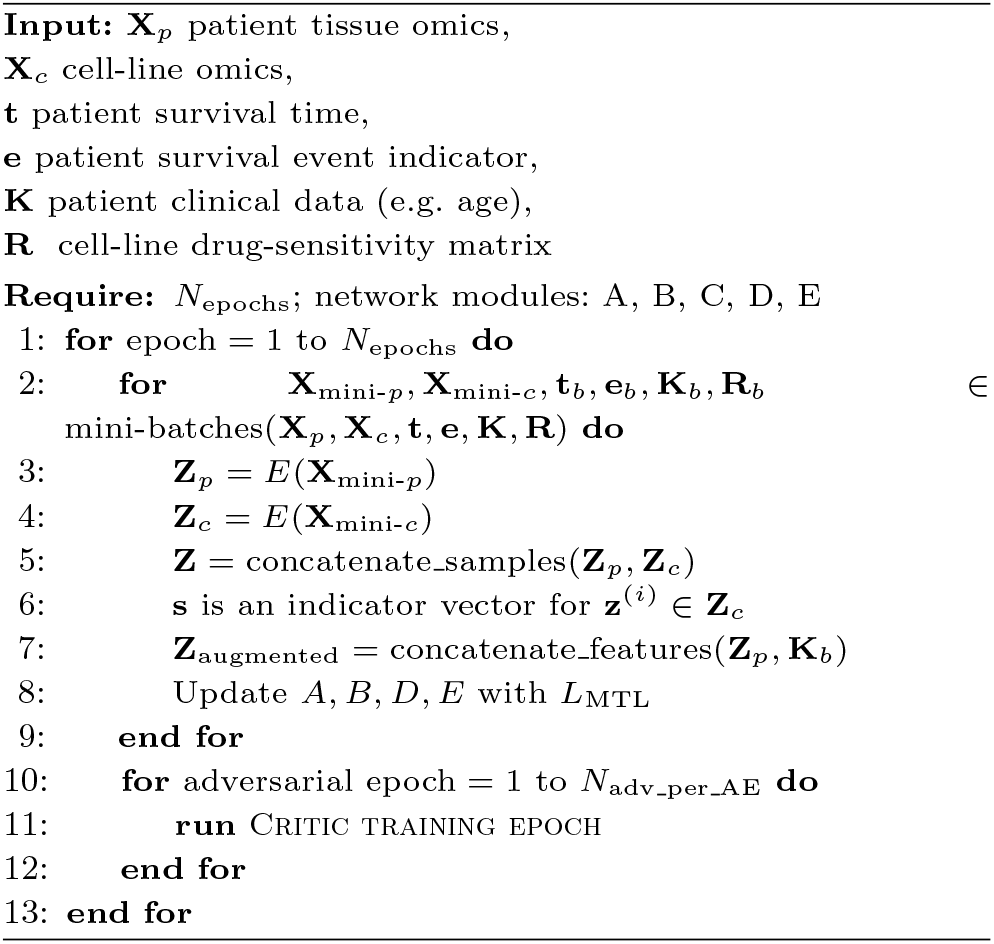

Survival data was obtained from Liu et al. (2018) who re-analysed TCGA clinical data to provide more accurate survival information. For drug sensitivity, we used CTRP which contains one of the most comprehensive drug-sensitivity datasets spanning 545 different drugs screened across 841 cell-lines of which 755 could be matched with transcriptomic profiles in CCLE (Seashore-Ludlow et al., 2015; Basu et al., 2013). CTRP data was downloaded using the *PharmacoGx* R-package (Smirnov et al., 2016). We decided to use the area above the curve (AAC) as the target drug-sensitivity value, since it is bounded in the range [0, 1] and is defined even when some other common alternatives, such as IC50, are not. Performance was evaluated with 5-fold cross-validation stratified with respect to the survival event and time in the case of patient data. Since the Cox model depends on the order of the events (Equation 6), event times were split into 10 equally sized quantiles after which the samples were evenly distributed into the folds based on the time quantile and event status. To account for the differences in time elapsed since TCGA and SCANB studies were conducted, the maximum follow-up time was set to 3000 days.

## Results

### Overview of MODAE

We developed MODAE to integrate and deconfound patient tissue and cell-line samples for survival and drug sensitivity related feature extraction in a hybrid unsupervised-supervised fashion (See Figure 1). Since survival and drug sensitivity are specific to their corresponding data domain, task specific models must be adapted to new target domains. MODAE achieves this by learning a shared representation of the molecular data which the task specific models can influence to mold it to be more useful for survival and drug sensitivity prediction. Differently from CODE-AE, we treat drug sensitivity prediction as a regression problem and perform multi-task learning with supervised tasks together with the reconstruction error.

**Fig. 1.**
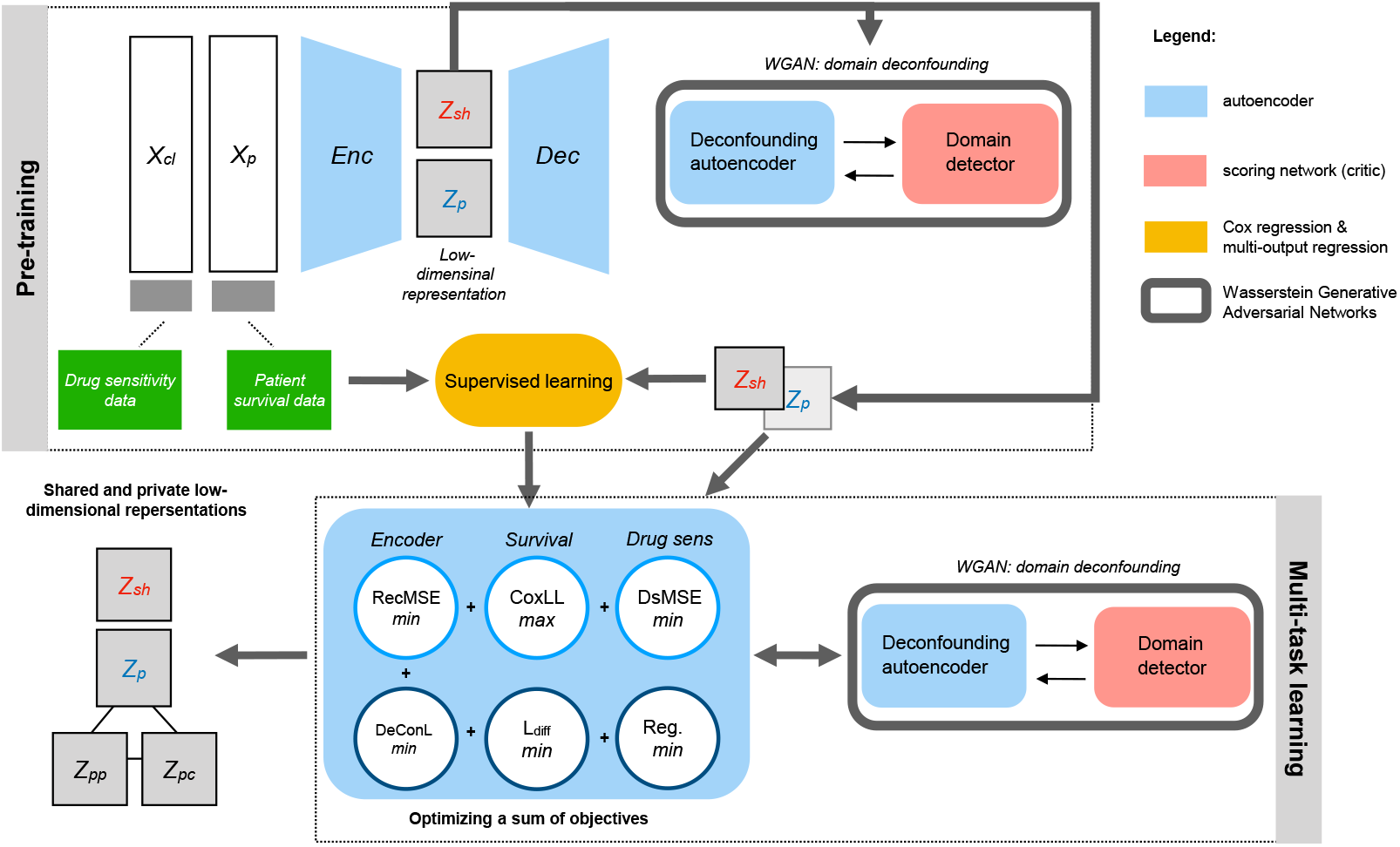
MODAE model diagram showing the essential structure and training steps. Cell-line omics profiles *X*_*cl*_ and patient omics profiles *X*_*p*_ are first encoded into two parts, the shared representation *z*_*sh*_ and private representation *z*_*p*_ which are further divided into private cell-line and private patient representations *z*_*pc*_ and *z*_*pp*_, respectively. The WGAN-based adversarial training adjusts the autoencoder with the help of a domain detector. Deconfounded shared representations are used for supervised training using drug-sensitivity and survival data. In a final training stage all tasks are trained simultaneously by optimizing a sum of objectives.

### Trade-offs between evaluation metrics in random hyper-parameter search

To search for the optimal hyper-parameters of MODAE, 100 iterations of random parameter search were performed. For the internal evaluation using 5-fold cross-validation in SCANB, the parameter search was carried out in a nested fashion by using one of the training folds as a validation set for parameter selection. Then for the test on external data (TCGA), the hyper-parameters were selected based on the 5-fold cross-validation results on SCANB. In either case, the selection of the best hyper-parameters is a multi-objective optimization problem. MODAE aims to optimize four different objectives: omics profile reconstruction, decondounding, survival prediction, and drug sensitivity prediction. The random parameter-search was intended to explore different trade-offs between these objectives by adjusting the learning rate, regularization, and number of hidden units associated with each module of MODAE as well as the weights assigned to the different objectives in Equation 8. Figure 2 shows the 5-fold average performance of each hyper-parameter setting: reconstruction error in negative *log*_10_*MSE* (separate for patient and cell-line data), deconfounding performance in terms of AUROC achieved by a linear SVM in distinguishing the shared embeddings corresponding to different datasets, survival concordance in Harrell’s C-index (Harrell, 1982), and 90^th^ percentile sensitivity prediction in *R*^2^. We evaluated each drug prediction separately and focus only on top 10% of drug sensitivity predictions because we wanted to focus on drugs for which the models could explain a large portion of variance across the cell-lines. Settings that were better in at least one metric compared to the others form a Pareto front which indicates results with the best trade-offs. The model corresponding to the highest sum of these metrics was compared to baseline models in cross-validation and external datasets.

**Fig. 2.**
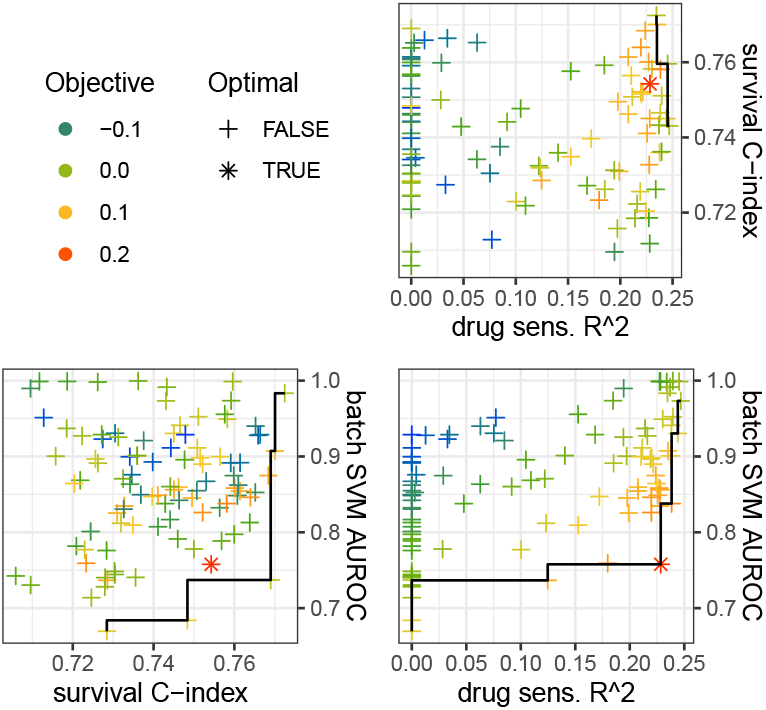
Pairwise scatter plots of key evaluation metrics. Each point represents one random parameter search result. The points are colored by the average sum of the test set survival concordance-index, top 10% drug sensitivity *R*^2^ quantile, and batch-effect defined as negative AUROC of a data domain detecting linear SVM. Drug-sensitivity performance in the plot was truncated at 0 since a portion of settings resulted in negative *R*^2^ suggesting a failure to learn useful models. The black line goes through points that are on the first Pareto front of each pairwise comparison. The optimal setting used for downstream analysis is highlighted by using an asterisk in place of a cross.

### Baseline models

As baseline, we used elastic net regression with simple feature selection and PCA. For drug-sensitivity prediction, each baseline drug model was trained independently and implemented with the *scikit-learn* Python module (Pedregosa et al., 2011). For survival we used the penalized Cox model implementation provided in the *scikit-survival* Python module (Pölsterl, 2020). The elastic net penalty and L1-ratio hyper-parameters were optimized via grid-search in nested cross-validation with the ranges [10^*−*3^, 10^*−*1^] and [0.1, 0.99], respectively. Feature selection was based on univariate regression models and the top-k approach, where the number of top predictors *k* was tuned as a hyper-parameter in the range [100, 1000]. These models were trained independently of each other and are therefore not considered multi-task learners, but their purpose is to provide context for the performance of MODAE.

### MODAE embeddings and performance comparison

Figures 3A-C show UMAP (McInnes et al., 2018) visualizations of the differences between cell-lines and patients in the original gene-expression, the deconfounded shared representation learned in pre-training, and the final shared representation learned while compromising between multiple tasks. The first plot shows a clear difference between the datasets with no sample from either dataset overlapping the other. Meanwhile the second plot demonstrates the power of the deconfounding autoencoder where the samples from CCLE evenly overlap the samples from SCANB. The third plot shows that the compromise with multi-task learning does not unravel the deconfounding effect learned during pre-training. Here all results are evaluated with the nested parameter search (i.e. the selected hyper-parameters can be different between folds). Figure 3D shows the survival concordance index of the models cross-validated in TCGA. All the models achieved concordance index around 0.75 with PCA based models performing better than univariate feature selection (UFS) and MODAE matching the best model in test set performance while showing slightly higher performance in the training set, but not to a large degree, which suggests that the model is adequately regularized to avoid overfitting. This result shows that MODAE achieves respectable performance even when burdened with deconfounding the predictive representation of differences between patient and cell-line profiles. Figure 3E shows the drug sensitivity prediction performance. Out of the three elastic net models, UFS performed the best, while 100 PCs was better than 10 PCs. MODAE performed significantly worse than the baselines, which suggests that the deconfounding process is hindering drug-sensitivity prediction. The 100 PC and UFS baseline models showed a significant degree of overfitting that was not present in MODAE. While the results shown here represent the average across best settings in each fold, it is consistent with Figure 2 which shows that no individual setting of MODAE performed significantly better than the search average.

**Fig. 3.**
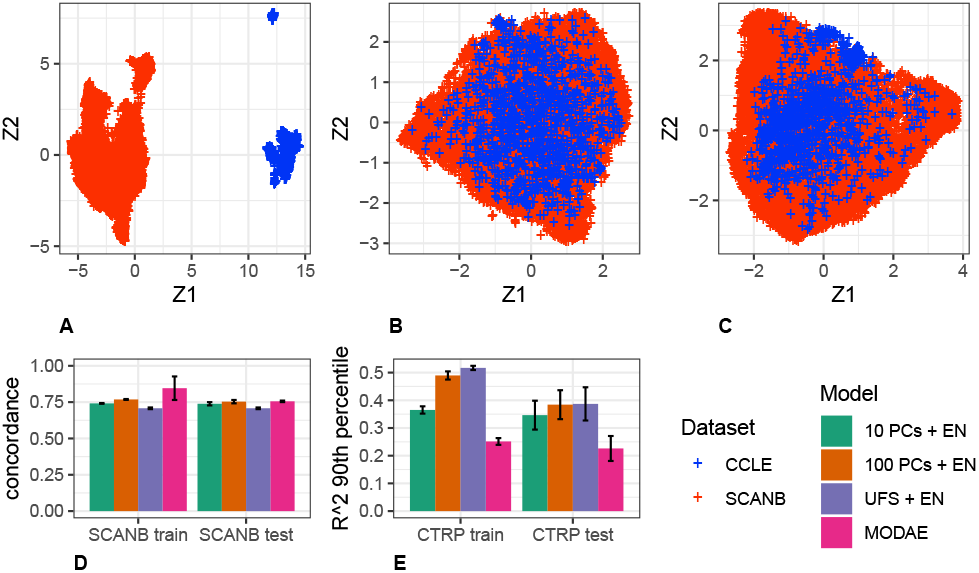
A) UMAP visualization of the concatenated standardized gene-expression matrices corresponding to SCANB and CCLE. **B)** UMAP visualization of the shared embeddings of SCANB and CCLE samples after pre-training the deconfounding autoencoder. **C)** UMAP visualization of the shared embeddings of SCANB and CCLE samples after joint multi-task learning. **D)** Mean survival concordance index of the proposed model and baselines cross-validated in SCANB. **E)** Mean of the 90^th^ percentile *R*^2^ of 545 drug sensitivity predictions cross validated in CTRP. Error bars represent standard deviation. PC: principal component; EN: elastic net regression; UFS: univariate feature selection; *R*^2^: coefficient of determination.

### Prediction performance on external survival dataset

We used the TCGA BRCA dataset as an external test set for survival predictions to see if the model predictions could be generalized to other datasets. The TCGA BRCA is a much smaller dataset but it has similar overall patient survival rate as the SCANB cohort with 16.3% in TCGA and 16.7% in SCANB. Since TCGA has much longer maximum follow up time, all observations beyond 3000 days were set to censored in both datasets to equalize both studies. Figure 4 shows the Harrell concordance index of MODAE as well as baseline models in the training set (SCANB) and the test set (TCGA). The external validation results reflect the cross-validation results with 100 PC EN and MODAE performing the best. The MODAE concordance index on the external dataset (0.710) was only slightly lower than on the internal test set (0.755) suggesting some losses in generalization. We also checked the proportional hazard assumption of MODAE by using the *survminer* -package in R. Supplementary Figure S2 shows the summary of tests on the last hidden layer of the survival risk module of MODAE, which indicated that the values observed at the hidden layers were not dependent on time and hence the proportional hazard assumption held for the final model. Due to differences in clinical data not all relevant covariates could be considered while testing the model. Additionally, the concordance index is based on the order of risk scores and does not account for shifts in predictions. Therefore, to further assess the predicted risk scores, we stratified the patients according to the predicted risk scores given by MODAE by calculating three even quantiles (high, mid, low) of the SCANB risk scores and stratifying TCGA patients using the same quantiles. Figure 5 shows the hazard ratios of a CoxPH model fitted with the additional clinical variables and the risk category in SCANB and Figure 6 shows the same for TCGA. In SCANB the group indicators, age z-score, and tumor size z-score were all significant while of the Nottingham histological grading only the third grade was significantly more predictive than the first grade. In TCGA only the high risk group from MODAE and the advanced tumor stages III and IV were significant compared to mid-risk and stage-I respectively. Notably the coefficients associated with the low-risk and high-risk groups were similar between the datasets, but the larger size of the SCANB cohort resulted in more significant p-values.

**Fig. 4.**
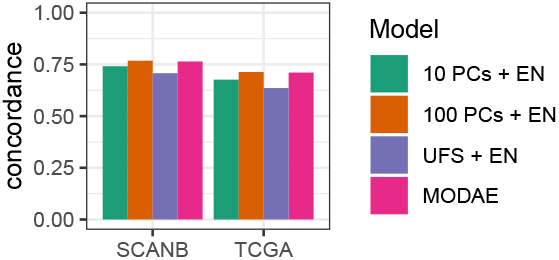
Survival prediction concordance index of MODAE and baseline models trained with SCANB and tested with TCGA BRCA.

**Fig. 5.**
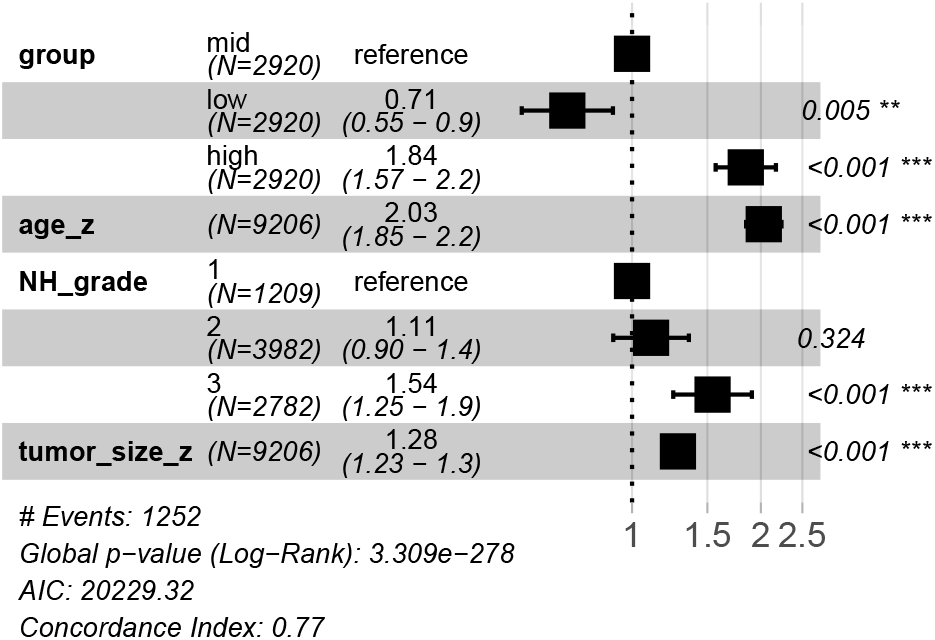
Hazard ratios of a CoxPH model with additional SCANB clinical variables and risk groups based on MODAE predictions.

**Fig. 6.**
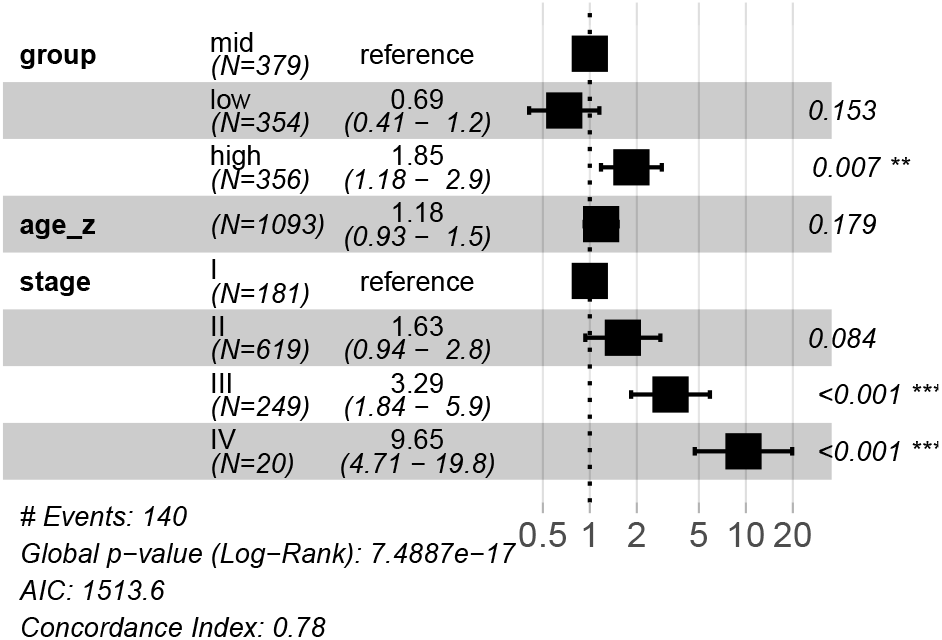
Hazard ratios of a CoxPH model with additional TCGA BRCA clinical variables and risk groups based on MODAE predictions.

### Assessing transfer learning of drug sensitivity predictions

The expression of the target genes modulated by a given drug is expected to be dysregulated. Similarly to He et al. (2022), to assess the agreement of the predictions with this expectation, we divided the patients into resistant and sensitive groups based on the sensitivity predicted by MODAE and compared the target-gene expression between them. The resistant and sensitive groups were defined as sensitivity z-score below *−*1 or above 1, respectively. 96% of gene-targets were dysregulated in breast cancer patient data with significance level 0.05, which suggests agreement with sensitivity predictions. However, only 22% of them had a fold-change over 1.5, which may suggest some false positives, since the sheer number of patients in SCANB can lend significance even to small differences. Supplementary File 1 contains the log-fold changes between groups and t-test p-values for every drug and their targets listed in DrugBank.

## Discussion

In this article we introduced a novel deep learning model to learn latent representations of omics data for predicting cancer patient survival and transfer of drug sensitivity predictions from cell-line screens to patients while adjusting for differences between the molecular profiles of these two sample types. We applied our model to a case study of breast cancer patients to illustrate its usefulness for precision medicine and patient stratification. The model performed well with survival prediction in cross-validation as well as external validation, suggesting that we succeeded in recapitulating clinically relevant patient information from their genomic profiles. However the drug sensitivity predictions on cell-lines were less accurate compared to simple linear models. A major limitation of this breast cancer case-study is the domain adaptation with pan-cancer cell-lines in the CCLE dataset. We wanted to focus on breast cancer, since the SCANB dataset is an unparalleled resource for studying a single cancer type and TCGA BRCA is the single largest cancer dataset within TCGA. Unfortunately there were not enough breast cancer cell-lines profiles available in CCLE to train reliable drug-sensitivity prediction models. In theory, the deconfounding autoencoder can learn to represent the other cancer types by using the private embeddings even if the other dataset only contains one type, however, the drug-sensitivity predictions are based on the shared embedding where cell-lines of different cancer types are all forced to a representation similar to breast cancer tissues. While the results are somewhat preliminary, they demonstrate that the model has the potential to aid in the discovery of new precision medicine therapies. In the future we aim to improve the drug-sensitivity model and extend the case-study to pancancer TCGA. Another interesting direction would be the use of drug-features (e.g., based on their chemical structures) since this could lead to targeted drug design for specific patient groups. Nevertheless, the proposed model could already be used to prioritize repositioning of drugs for high-risk patients.

## Competing interests

The authors declare no competing interests.

## Author contributions

T.J.R. and V.F. conceived the survival-drug-sensitivity multitask model; F.N. contributed to the adversarial model conceptualization; T.J.R. developed the methods, software, and experiments; T.J.R. wrote the first manuscript draft and revised it together with V.F.; V.F. and F.N. supervised the project; V.F. acquired the main grant while T.J.R. obtained an additional grant to support the project. All authors have read and agreed to the published version of the manuscript.

## Acknowledgements

Computations were performed on infrastructure hosted by the Bioinformatics Center, University of Eastern Finland and the Finnish IT center for science (CSC).

## Funding

This work was supported by the Academy of Finland [grant numbers 336275, 332510], the Jane and Aatos Erkko Foundation, Sigrid Jusélius Foundation, and the Finnish Cultural Foundation [grant number 00230994 to T.J.R.].

## Software availability

https://github.com/UEFBiomedicalInformaticsLab/MODAE

## Notes

### Competing Interest Statement

The authors have declared no competing interest.

### Summary of Updates

Result section on objective trade-offs updated to clarify parameter search validation strategy. Author contributions updated to be more precise.

