## Supplementary Figures and Tables for "Multi-task deep latent spaces for cancer survival and drug sensitivity prediction"

### Supplementary algorithms, figures, and tables for: Multi-task deep latent spaces for cancer survival and treatment response prediction

Teemu J. Rintala [1], Francesco Napolitano [2], and Vittorio Fortino [1,*]

[1] Institute of Biomedicine, University of Eastern Finland, 70210 Kuopio, Finland
[2] Department of Science and Technology, University of Sannio, 82100 Benevento, Italy
[*] To whom correspondence should be addressed:


#### 1 Supplementary Algorithms

**Algorithm 1** Pre-training epochs

**Input:** $\mathbf{X}_p$ patient tissue omics,
$\mathbf{X}_c$ cell-line omics,
$\mathbf{t}$ patient survival time,
$\mathbf{e}$ patient survival event indicator,
$\mathbf{K}$ patient clinical data (e.g., age),
$\mathbf{R}$ cell-line drug-sensitivity matrix

**Require:** $N_{\text{adv\_per\_AE}}$; network modules: A, B, C, D, E

**procedure** AUTOENCODER PRE-TRAINING EPOCH
    **for** $\mathbf{X}_{\text{mini-}p}, \mathbf{X}_{\text{mini-}c} \in \text{minibatches}(\mathbf{X}_p, \mathbf{X}_c)$ **do**
        $\mathbf{X}_{\text{mini}} = \text{concatenate\_samples}(\mathbf{X}_{\text{mini-}p}, \mathbf{X}_{\text{mini-}c})$
        Update $E$ and $D$ with $L_{\text{rec}}$
    **end for**
**end procedure**
**procedure** CRITIC TRAINING EPOCH
    **for** $\mathbf{X}_{\text{mini-}p}, \mathbf{X}_{\text{mini-}c} \in \text{minibatches}(\mathbf{X}_p, \mathbf{X}_c)$ **do**
        $\mathbf{X}_{\text{mini}} = \text{concatenate\_samples}(\mathbf{X}_{\text{mini-}p}, \mathbf{X}_{\text{mini-}c})$
        $\mathbf{s}$ is an indicator vector for $\mathbf{x}^{(i)} \in \mathbf{X}_{\text{mini-}c}$
        $\mathbf{Z} = E(\mathbf{X}_{\text{mini}})$
        Update $C$ with $L_{\text{adv}}(\mathbf{Z}, \mathbf{s})$
    **end for**
**end procedure**
**procedure** BATCH CORRECTION PRE-TRAINING EPOCH
    **for** $\mathbf{X}_{\text{mini-}p}, \mathbf{X}_{\text{mini-}c} \in \text{minibatches}(\mathbf{X}_p, \mathbf{X}_c)$ **do**
        $\mathbf{X}_{\text{mini}} = \text{concatenate\_samples}(\mathbf{X}_{\text{mini-}p}, \mathbf{X}_{\text{mini-}c})$
        $\mathbf{s}$ is an indicator vector for $\mathbf{x}^{(i)} \in \mathbf{X}_{\text{mini-}c}$
        $\mathbf{Z} = E(\mathbf{X}_{\text{mini}})$
        Update $C$ with $L_{\text{bc}}(\mathbf{Z}, \mathbf{s})$
    **end for**
    **for** adversarial epoch = 1 to $N_{\text{adv\_per\_AE}}$ **do**
        **run** CRITIC TRAINING EPOCH
    **end for**
**end procedure**
**procedure** SURVIVAL PRE-TRAINING EPOCH
    **for** $\mathbf{X}_{\text{mini}}, \mathbf{t}_b, \mathbf{e}_b, \mathbf{K}_b \in \text{minibatches}(\mathbf{X}_p, \mathbf{t}, \mathbf{e}, \mathbf{K})$ **do**
        $\mathbf{Z} = E(\mathbf{X}_{\text{mini}})$
        $\mathbf{Z}_{\text{augmented}} = \mathbf{Z} \oplus \mathbf{K}_b$
        Update $A$ with $L_{\text{surv}}$
    **end for**
**end procedure**
**procedure** DRUG SENSITIVITY PRE-TRAINING EPOCH
    **for** $\mathbf{X}_{\text{mini}}, \mathbf{R}_b \in \text{minibatches}(\mathbf{X}_c, \mathbf{R})$ **do**
        $\mathbf{Z} = E(\mathbf{X}_{\text{mini}})$
        Update $B$ with $L_{\text{drug}}$
    **end for**
**end procedure**

**Algorithm 2** Pre-training network modules

**Require:** $N_{\text{AE-pte}}, N_{\text{C-pte}}, N_{\text{BC-pte}}, N_{\text{S-pte}}, N_{\text{D-pte}}$
**Require:** network modules: A, B, C, D, E

```
for epoch = 1 to N_AE-pte do
    run Autoencoder pre-training epoch
end for
for epoch = 1 to N_C-pte do
    run Critic training epoch
end for
for epoch = 1 to N_BC-pte do
    run Batch correction pre-training epoch
end for
for epoch = 1 to N_S-pte do
    run Survival pre-training epoch
end for
for epoch = 1 to N_D-pte do
    run Drug sensitivity pre-training epoch
end for
```

### 2 Supplementary figures and tables

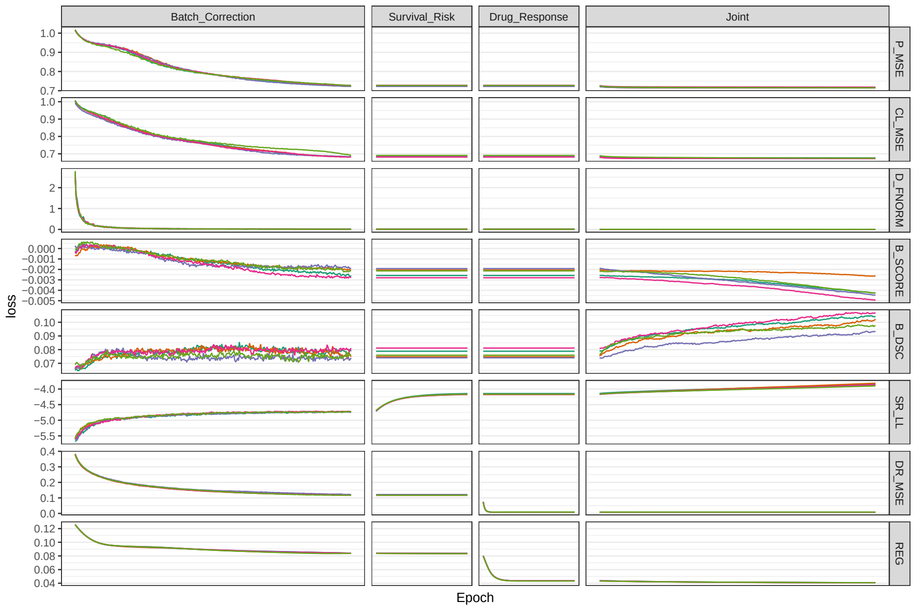


Figure 1: MODAE losses and additional metrics monitored over epochs. Different color lines represent different cross-validation folds. P_MSE = patient omic reconstruction mean squared error; CL_MSE = cell-line omic reconstruction mean squared error; D_FNORM = deconfounding norm penalty; B_SCORE = average critic score difference between patient and cell-line representations (corresponds to $L_{adv}$); B_DSC = dispersion separability criterion between patients and cell-line representations; SR_LL = Cox PH log-likelihood (higher is better); DR_MSE = drug-sensitivity regression mean squared error; REG = total losses from L2-norm penalty on model weights.

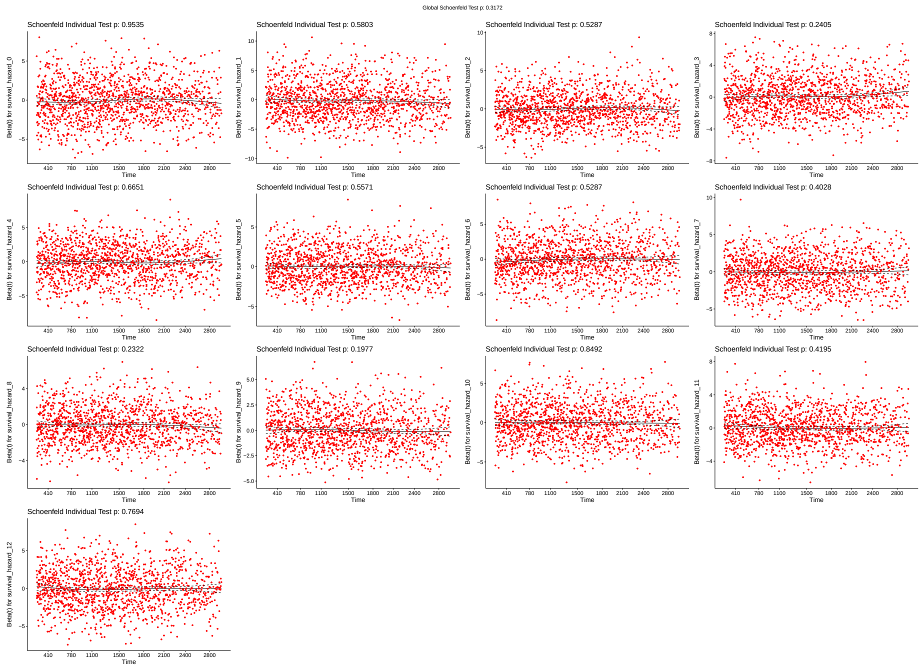


Figure 2: MODAE survival model Cox PH assumption tests based on last hidden layer values. No hidden unit was significantly associated with survival time, therefore the proportional hazards assumption holds.

Table 1: Gene-expression difference breast cancer patients stratified based on drug sensitivity. Genes identified of breast cancer drug targets. P-value was determined using the t-test.

| drug | gene | log2FC | p_value |
|---|---|---|---|
| Abiraterone | CYP17A1 | -0.015 | 0.001 |
| Afatinib | EGFR | 0.733 | <0.001 |
| Afatinib | ERBB2 | -0.579 | <0.001 |
| Afatinib | ERBB4 | -1.371 | <0.001 |
| Alisertib | AURKA | -1.154 | <0.001 |
| Alvocidib | EGFR | -0.704 | <0.001 |
| Alvocidib | PYGB | 0.335 | <0.001 |
| Alvocidib | PYGM | 0.072 | 0.001 |
| Alvocidib | CDK7 | -0.050 | 0.005 |
| Alvocidib | CDK5 | 0.255 | <0.001 |
| Alvocidib | CDK9 | 0.286 | <0.001 |
| Alvocidib | CDK1 | -1.084 | <0.001 |
| Alvocidib | CDK6 | -0.665 | <0.001 |

| drug | gene | log2FC | p-value |
|---|---|---|---|
| Alvocidib | CDK4 | -0.242 | <0.001 |
| Alvocidib | CDK8 | -0.484 | <0.001 |
| Alvocidib | CDK2 | -0.302 | <0.001 |
| Axitinib | FLT1 | -0.115 | <0.001 |
| Axitinib | KDR | -0.058 | 0.019 |
| Axitinib | FLT4 | 0.457 | <0.001 |
| Azacitidine | DNMT1 | -0.343 | <0.001 |
| Bardoxolone methyl | NFKBIA | -0.116 | <0.001 |
| Belinostat | HDAC1 | -0.174 | <0.001 |
| Bexarotene | RXRA | 0.955 | <0.001 |
| Bexarotene | RXRB | 0.175 | <0.001 |
| Bexarotene | RXRG | 0.058 | <0.001 |
| Bortezomib | PSMB5 | -0.127 | <0.001 |
| Bortezomib | PSMB1 | -0.287 | <0.001 |
| Bosutinib | CDK2 | -0.303 | <0.001 |
| Bosutinib | BCR | 0.185 | <0.001 |
| Bosutinib | ABL1 | 0.348 | <0.001 |
| Bosutinib | LYN | -0.819 | <0.001 |
| Bosutinib | HCK | -0.357 | <0.001 |
| Bosutinib | SRC | 0.133 | <0.001 |
| Bosutinib | MAP2K1 | -0.143 | <0.001 |
| Bosutinib | MAP2K2 | 0.382 | <0.001 |
| Bosutinib | MAP3K2 | -0.080 | <0.001 |
| Bosutinib | CAMK2G | -0.101 | <0.001 |
| Cabozantinib | KDR | -0.055 | 0.024 |
| Cabozantinib | MET | -0.623 | <0.001 |
| Cabozantinib | RET | 1.355 | <0.001 |
| Canertinib | EGFR | -0.698 | <0.001 |
| Cediranib | KDR | -0.056 | 0.023 |
| Cerulenin | FASN | 1.390 | <0.001 |
| Ciclopirox | ATP1A1 | -0.037 | 0.05 |
| Ciclosporin | CAMLG | 0.281 | <0.001 |
| Ciclosporin | PPP3R2 | -0.000 | 0.736 |
| Ciclosporin | PPIA | -0.346 | <0.001 |
| Ciclosporin | PPIF | -0.189 | <0.001 |
| Cimetidine | HRH2 | 0.211 | <0.001 |
| Clofarabine | POLA1 | -0.303 | <0.001 |
| Clofarabine | RRM1 | -0.279 | <0.001 |
| Crizotinib | MET | -0.623 | <0.001 |
| Crizotinib | ALK | -0.140 | <0.001 |
| Crizotinib | ROS1 | -0.131 | <0.001 |
| Crizotinib | MST1R | 0.771 | <0.001 |
| Cyclophosphamide | NR1I2 | -0.064 | <0.001 |
| Dabrafenib | BRAF | -0.085 | <0.001 |
| Dabrafenib | RAF1 | -0.123 | <0.001 |

| drug | gene | log2FC | p-value |
|---|---|---|---|
| Dabrafenib | SIK1 | 0.395 | <0.001 |
| Dabrafenib | NEK11 | 0.600 | <0.001 |
| Dabrafenib | LIMK1 | 0.260 | <0.001 |
| Dacarbazine | POLA2 | -0.410 | <0.001 |
| Dacarbazine | PGD | -0.092 | <0.001 |
| Daporinad | NAMPT | -0.337 | <0.001 |
| Dasatinib | BCR | -0.187 | <0.001 |
| Dasatinib | ABL1 | -0.330 | <0.001 |
| Dasatinib | LYN | 0.872 | <0.001 |
| Dasatinib | SRC | -0.136 | <0.001 |
| Dasatinib | EPHA2 | -0.154 | <0.001 |
| Dasatinib | LCK | 1.237 | <0.001 |
| Dasatinib | YES1 | 0.422 | <0.001 |
| Dasatinib | KIT | 0.426 | <0.001 |
| Dasatinib | PDGFRB | -0.428 | <0.001 |
| Dasatinib | STAT5B | -0.122 | <0.001 |
| Dasatinib | ABL2 | 0.104 | <0.001 |
| Dasatinib | FYN | 0.730 | <0.001 |
| Dasatinib | BTK | 0.730 | <0.001 |
| Dasatinib | NR4A3 | 0.321 | <0.001 |
| Dasatinib | CSK | -0.086 | 0.005 |
| Dasatinib | EPHA5 | -0.003 | 0.585 |
| Dasatinib | EPHB4 | -0.281 | <0.001 |
| Dasatinib | FGR | 0.390 | <0.001 |
| Dasatinib | FRK | -0.108 | <0.001 |
| Dasatinib | HSPA8 | 0.262 | <0.001 |
| Dasatinib | MAP3K20 | -0.142 | <0.001 |
| Dasatinib | MAPK14 | 0.245 | <0.001 |
| Dasatinib | PPAT | 0.345 | <0.001 |
| Decitabine | DNMT1 | -0.342 | <0.001 |
| Decitabine | HDAC1 | -0.174 | <0.001 |
| Decitabine | DNMT3A | -0.174 | <0.001 |
| Decitabine | DNMT3B | -0.789 | <0.001 |
| Dexamethasone | NR1I2 | -0.064 | <0.001 |
| Dexamethasone | NR3C1 | -0.117 | <0.001 |
| Dexamethasone | NR0B1 | -0.033 | 0.056 |
| Dexamethasone | ANXA1 | -0.543 | <0.001 |
| Dexamethasone | NOS2 | -0.018 | 0.21 |
| Docetaxel | NR1I2 | -0.065 | <0.001 |
| Docetaxel | TUBB1 | 0.018 | 0.067 |
| Docetaxel | MAP2 | -0.451 | <0.001 |
| Docetaxel | MAP4 | 0.156 | <0.001 |
| Docetaxel | MAPT | 2.135 | <0.001 |
| Docetaxel | BCL2 | 0.840 | <0.001 |
| Doxorubicin | TOP2A | -1.180 | <0.001 |

| drug | gene | log2FC | p-value |
|---|---|---|---|
| Doxorubicin | NOLC1 | -0.167 | <0.001 |
| Doxorubicin | TOP1 | -0.275 | <0.001 |
| Doxorubicin | TOP2B | -0.162 | <0.001 |
| Elocalcitol | VDR | 0.241 | <0.001 |
| Entinostat | HDAC1 | -0.173 | <0.001 |
| Erismodegib | SMO | 0.074 | 0.036 |
| Erlotinib | EGFR | 0.718 | <0.001 |
| Erlotinib | NR1I2 | 0.064 | <0.001 |
| Etoposide | TOP2A | -1.178 | <0.001 |
| Etoposide | TOP2B | -0.161 | <0.001 |
| Fingolimod | HDAC1 | -0.174 | <0.001 |
| Fingolimod | S1PR5 | -0.111 | <0.001 |
| Fingolimod | S1PR1 | -0.043 | 0.183 |
| Fingolimod | S1PR3 | 0.048 | <0.001 |
| Fingolimod | S1PR4 | -0.469 | <0.001 |
| Fluorouracil | TYMS | -0.919 | <0.001 |
| Fluvastatin | HMGCR | -0.084 | <0.001 |
| Fluvastatin | HDAC2 | -0.904 | <0.001 |
| Foretinib | KDR | -0.055 | 0.024 |
| Foretinib | HGF | 0.077 | 0.014 |
| Fulvestrant | ESR1 | 2.923 | <0.001 |
| Gefitinib | EGFR | 0.728 | <0.001 |
| Gemcitabine | RRM1 | -0.279 | <0.001 |
| Gemcitabine | TYMS | -0.919 | <0.001 |
| Gemcitabine | CMPK1 | -0.311 | <0.001 |
| Ibrutinib | BTK | -0.682 | <0.001 |
| Ifosfamide | NR1I2 | 0.064 | <0.001 |
| Imatinib | BCR | 0.188 | <0.001 |
| Imatinib | ABL1 | 0.347 | <0.001 |
| Imatinib | KIT | -0.384 | <0.001 |
| Imatinib | PDGFRB | 0.515 | <0.001 |
| Imatinib | NTRK1 | 0.155 | <0.001 |
| Imatinib | CSF1R | 0.123 | <0.001 |
| Imatinib | PDGFRA | -0.322 | <0.001 |
| Imatinib | DDR1 | 0.588 | <0.001 |
| Imatinib | DDR2 | -0.134 | <0.001 |
| Istradefylline | ADORA2A | 0.163 | <0.001 |
| Istradefylline | ADORA1 | 0.263 | <0.001 |
| Itraconazole | CYP51A1 | -0.031 | 0.165 |
| Lapatinib | EGFR | -0.703 | <0.001 |
| Lapatinib | ERBB2 | 0.648 | <0.001 |
| Lenvatinib | FLT1 | -0.112 | <0.001 |
| Lenvatinib | KDR | -0.056 | 0.023 |
| Lenvatinib | FLT4 | 0.459 | <0.001 |
| Lenvatinib | RET | 1.353 | <0.001 |

| drug | gene | log2FC | p-value |
|---|---|---|---|
| Lenvatinib | KIT | -0.383 | <0.001 |
| Lenvatinib | PDGFRA | -0.322 | <0.001 |
| Lenvatinib | FGFR1 | 0.229 | <0.001 |
| Lenvatinib | FGFR2 | 0.487 | <0.001 |
| Lenvatinib | FGFR3 | 0.799 | <0.001 |
| Lenvatinib | FGFR4 | -0.202 | <0.001 |
| Linifanib | FLT1 | -0.114 | <0.001 |
| Linifanib | KDR | -0.057 | 0.021 |
| Linifanib | FLT4 | 0.457 | <0.001 |
| Linifanib | KIT | -0.377 | <0.001 |
| Linifanib | CSF1R | 0.124 | <0.001 |
| Linifanib | FLT3 | 0.136 | 0.004 |
| Linsitinib | INSR | 0.326 | <0.001 |
| Linsitinib | IGF1R | 1.342 | <0.001 |
| Lovastatin | HMGCR | -0.090 | <0.001 |
| Lovastatin | HDAC2 | -0.912 | <0.001 |
| Lovastatin | ITGAL | -0.567 | <0.001 |
| Methotrexate | TYMS | -0.921 | <0.001 |
| Methotrexate | ATIC | -0.053 | <0.001 |
| Methotrexate | DHFR | -0.370 | <0.001 |
| Myricetin | JAK1 | -0.072 | <0.001 |
| Myricetin | PIK3CG | -0.548 | <0.001 |
| Navitoclax | BCL2 | 0.840 | <0.001 |
| Navitoclax | BCL2L2 | 0.142 | <0.001 |
| Navitoclax | BCL2L2-PABPN1 | 0.243 | <0.001 |
| Navitoclax | BAD | 0.868 | <0.001 |
| Nelarabine | POLA1 | -0.303 | <0.001 |
| Neratinib | EGFR | -0.699 | <0.001 |
| Nilotinib | ABL1 | 0.347 | <0.001 |
| Nilotinib | KIT | -0.384 | <0.001 |
| Nintedanib | FLT1 | -0.112 | <0.001 |
| Nintedanib | KDR | -0.056 | 0.023 |
| Nintedanib | FLT4 | 0.459 | <0.001 |
| Nintedanib | LYN | -0.818 | <0.001 |
| Nintedanib | SRC | 0.133 | <0.001 |
| Nintedanib | LCK | -1.197 | <0.001 |
| Nintedanib | PDGFRB | 0.515 | <0.001 |
| Nintedanib | PDGFRA | -0.320 | <0.001 |
| Nintedanib | FGFR1 | 0.229 | <0.001 |
| Nintedanib | FGFR2 | 0.489 | <0.001 |
| Nintedanib | FGFR3 | 0.799 | <0.001 |
| Nintedanib | FLT3 | 0.139 | 0.003 |
| Obatoclax | BCL2 | 0.840 | <0.001 |
| Olaparib | PARP1 | -0.127 | <0.001 |
| Olaparib | PARP2 | -0.380 | <0.001 |

| drug | gene | log2FC | p-value |
|---|---|---|---|
| Olaparib | PARP3 | 0.665 | <0.001 |
| Olaparib | AKR1C3 | 0.133 | 0.003 |
| Omacetaxine mepesuccinate | RPL3 | 0.317 | <0.001 |
| Paclitaxel | NR1I2 | -0.065 | <0.001 |
| Paclitaxel | TUBB1 | 0.018 | 0.067 |
| Paclitaxel | MAP2 | -0.451 | <0.001 |
| Paclitaxel | MAP4 | 0.155 | <0.001 |
| Paclitaxel | MAPT | 2.135 | <0.001 |
| Paclitaxel | BCL2 | 0.841 | <0.001 |
| Pazopanib | FLT1 | -0.115 | <0.001 |
| Pazopanib | KDR | -0.058 | 0.019 |
| Pazopanib | FLT4 | 0.458 | <0.001 |
| Pazopanib | KIT | -0.379 | <0.001 |
| Pazopanib | PDGFRB | 0.515 | <0.001 |
| Pazopanib | PDGFRA | -0.319 | <0.001 |
| Pazopanib | FGFR3 | 0.798 | <0.001 |
| Pazopanib | ITK | -1.238 | <0.001 |
| Pazopanib | FGF1 | 0.284 | <0.001 |
| Pazopanib | SH2B3 | -0.227 | <0.001 |
| Procarbazine | MAOA | 0.074 | 0.284 |
| Prochlorperazine | DRD2 | 0.053 | 0.024 |
| Prochlorperazine | HRH1 | 0.017 | 0.584 |
| Prochlorperazine | ADRA1A | 0.023 | 0.237 |
| Prochlorperazine | ADRA2A | 0.857 | <0.001 |
| Quizartinib | FLT3 | 0.137 | 0.004 |
| Regorafenib | FLT1 | -0.114 | <0.001 |
| Regorafenib | KDR | -0.057 | 0.02 |
| Regorafenib | FLT4 | 0.458 | <0.001 |
| Regorafenib | ABL1 | 0.347 | <0.001 |
| Regorafenib | RET | 1.349 | <0.001 |
| Regorafenib | BRAF | -0.086 | <0.001 |
| Regorafenib | RAF1 | -0.123 | <0.001 |
| Regorafenib | EPHA2 | 0.184 | <0.001 |
| Regorafenib | KIT | -0.378 | <0.001 |
| Regorafenib | PDGFRB | 0.516 | <0.001 |
| Regorafenib | FRK | 0.130 | <0.001 |
| Regorafenib | NTRK1 | 0.155 | <0.001 |
| Regorafenib | PDGFRA | -0.318 | <0.001 |
| Regorafenib | DDR2 | -0.132 | <0.001 |
| Regorafenib | FGFR1 | 0.230 | <0.001 |
| Regorafenib | FGFR2 | 0.490 | <0.001 |
| Regorafenib | TEK | 0.113 | <0.001 |
| Regorafenib | MAPK11 | 0.377 | <0.001 |
| Ruxolitinib | JAK1 | -0.072 | <0.001 |
| Ruxolitinib | JAK2 | -0.382 | <0.001 |

| drug | gene | log2FC | p-value |
|---|---|---|---|
| Ruxolitinib | JAK3 | -0.825 | <0.001 |
| Ruxolitinib | TYK2 | 0.225 | <0.001 |
| Selumetinib | MAP2K1 | 0.142 | <0.001 |
| Selumetinib | MAP2K2 | -0.384 | <0.001 |
| Sildenafil | PDE5A | 0.003 | 0.941 |
| Sildenafil | PDE6G | -0.439 | <0.001 |
| Sildenafil | PDE6H | -0.012 | 0.011 |
| Sildenafil | ODC1 | -0.788 | <0.001 |
| Sildenafil | CD274 | -0.952 | <0.001 |
| Simvastatin | HMGCR | 0.088 | <0.001 |
| Simvastatin | HDAC2 | 0.883 | <0.001 |
| Simvastatin | ITGAL | 0.571 | <0.001 |
| Sirolimus | MTOR | -0.040 | <0.001 |
| Sitagliptin | DPP4 | 0.322 | <0.001 |
| Sorafenib | FLT1 | -0.115 | <0.001 |
| Sorafenib | KDR | -0.057 | 0.02 |
| Sorafenib | FLT4 | 0.458 | <0.001 |
| Sorafenib | RET | 1.351 | <0.001 |
| Sorafenib | BRAF | -0.086 | <0.001 |
| Sorafenib | RAF1 | -0.123 | <0.001 |
| Sorafenib | KIT | -0.379 | <0.001 |
| Sorafenib | PDGFRB | 0.516 | <0.001 |
| Sorafenib | FGFR1 | 0.229 | <0.001 |
| Sorafenib | FLT3 | 0.138 | 0.003 |
| Sunitinib | FLT1 | -0.113 | <0.001 |
| Sunitinib | KDR | -0.055 | 0.024 |
| Sunitinib | FLT4 | 0.459 | <0.001 |
| Sunitinib | MET | -0.622 | <0.001 |
| Sunitinib | KIT | -0.377 | <0.001 |
| Sunitinib | PDGFRB | 0.516 | <0.001 |
| Sunitinib | CSF1R | 0.125 | <0.001 |
| Sunitinib | PDGFRA | -0.316 | <0.001 |
| Sunitinib | FLT3 | 0.138 | 0.003 |
| Tacrolimus | FKBP1A | -0.457 | <0.001 |
| Tamoxifen | NR1I2 | -0.064 | <0.001 |
| Tamoxifen | ESR1 | 2.927 | <0.001 |
| Tamoxifen | ESR2 | -0.129 | <0.001 |
| Tamoxifen | PRKCA | -0.353 | <0.001 |
| Tamoxifen | SHBG | 0.065 | <0.001 |
| Tamoxifen | EBP | -0.181 | <0.001 |
| Tamoxifen | AR | 0.964 | <0.001 |
| Tamoxifen | KCNH2 | 0.127 | 0.001 |
| Tamoxifen | ESRRG | 0.278 | <0.001 |
| Tamoxifen | MAPK8 | -0.148 | <0.001 |
| Tandutinib | FLT3 | 0.137 | 0.003 |

| drug | gene | log2FC | p-value |
|---|---|---|---|
| Tandutinib | PDGFD | 0.590 | <0.001 |
| Tanespimycin | HSP90AA1 | -0.381 | <0.001 |
| Tanespimycin | HSP90AB1 | -0.301 | <0.001 |
| Temsirolimus | MTOR | -0.040 | <0.001 |
| Teniposide | TOP2A | -1.177 | <0.001 |
| Thalidomide | CRBN | 0.074 | <0.001 |
| Thalidomide | ORM1 | 0.452 | <0.001 |
| Tivantinib | MET | -0.625 | <0.001 |
| Tivozanib | FLT1 | -0.111 | <0.001 |
| Tivozanib | KDR | -0.056 | 0.022 |
| Tivozanib | FLT4 | 0.458 | <0.001 |
| Tivozanib | MET | -0.626 | <0.001 |
| Tivozanib | KIT | -0.384 | <0.001 |
| Tivozanib | PDGFRB | 0.515 | <0.001 |
| Tivozanib | PDGFRA | -0.322 | <0.001 |
| Tivozanib | FGFR1 | 0.228 | <0.001 |
| Tivozanib | FLT3 | 0.137 | 0.004 |
| Tivozanib | TEK | 0.113 | <0.001 |
| Tivozanib | PTK6 | 0.911 | <0.001 |
| Topotecan | TOP1 | -0.275 | <0.001 |
| Topotecan | TOP1MT | -0.058 | 0.017 |
| Tosedostat | NPEPPS | 0.194 | <0.001 |
| Tosedostat | LTA4H | -0.262 | <0.001 |
| Trametinib | MAP2K1 | 0.147 | <0.001 |
| Trametinib | MAP2K2 | -0.388 | <0.001 |
| Tretinoin | RXRA | 0.956 | <0.001 |
| Tretinoin | RXRB | 0.175 | <0.001 |
| Tretinoin | RXRG | 0.058 | <0.001 |
| Tretinoin | RARG | 0.390 | <0.001 |
| Tretinoin | ALDH1A1 | -0.324 | <0.001 |
| Tretinoin | GPRC5A | 0.890 | <0.001 |
| Tretinoin | ALDH1A2 | 0.216 | <0.001 |
| Tretinoin | RARRES1 | -1.769 | <0.001 |
| Tretinoin | RARA | 1.091 | <0.001 |
| Tretinoin | RARB | -0.512 | <0.001 |
| Tretinoin | LCN1 | 0.011 | 0.034 |
| Tretinoin | OBP2A | 0.399 | <0.001 |
| Tretinoin | RBP4 | 0.337 | <0.001 |
| Tretinoin | PDK4 | 0.168 | 0.004 |
| Tretinoin | CYP26A1 | 0.294 | <0.001 |
| Tretinoin | CYP26B1 | -0.153 | <0.001 |
| Tretinoin | CYP26C1 | 0.013 | <0.001 |
| Tretinoin | HPGDS | 0.395 | <0.001 |
| Trifluoperazine | DRD2 | 0.054 | 0.023 |
| Trifluoperazine | ADRA1A | 0.023 | 0.229 |

| drug | gene | log2FC | p-value |
|---|---|---|---|
| Trifluoperazine | CALY | 0.125 | <0.001 |
| Trifluoperazine | CALM1 | -0.271 | <0.001 |
| Trifluoperazine | TNNC1 | 0.353 | <0.001 |
| Trifluoperazine | S100A4 | -0.395 | <0.001 |
| Valdecoxib | PTGS2 | -0.352 | <0.001 |
| Valdecoxib | CA2 | -0.019 | 0.777 |
| Valdecoxib | CA3 | -0.105 | 0.018 |
| Vandetanib | EGFR | 0.725 | <0.001 |
| Vandetanib | RET | -1.358 | <0.001 |
| Vandetanib | TEK | -0.105 | <0.001 |
| Vandetanib | PTK6 | -0.915 | <0.001 |
| Vandetanib | VEGFA | 0.069 | 0.026 |
| Veliparib | PARP1 | -0.125 | <0.001 |
| Veliparib | PARP2 | -0.379 | <0.001 |
| Vincristine | TUBB | -0.360 | <0.001 |
| Vincristine | TUBA4A | -0.378 | <0.001 |
| Vorapaxar | F2R | 0.072 | 0.019 |
| Vorinostat | HDAC1 | -0.174 | <0.001 |
| Vorinostat | HDAC2 | -0.903 | <0.001 |
| Vorinostat | HDAC3 | -0.031 | 0.002 |
| Vorinostat | HDAC6 | 0.164 | <0.001 |
| Vorinostat | HDAC8 | 0.018 | 0.142 |
